# Beyond viability: Seed ageing alters development and phenology of adult plants

**DOI:** 10.64898/2026.04.23.720367

**Authors:** Lea Klepka, Sascha Liepelt, Paola Ariadna Calles Monzón, Susanna Konrad, Anna Bucharova

**Affiliations:** Conservation Biology, Department of Biology, Philipps University Marburg

**Keywords:** artificial seed ageing, multi-species common garden, conservation seed bank, direct storage effects, invisible fraction, life-history traits, seed deterioration

## Abstract

1. Stored seeds are crucial repositories of plant genetic diversity. However, long-term storage inevitably causes seed deterioration and loss of viability, and chemical processes within the seeds during storage can influence germination and seedling establishment. Emerging evidence suggests that seed ageing can also affect traits of adult plants, yet the extent to which this phenomenon is relevant across species, particularly for wild plant species with high genetic variation, remains unclear.
2. To address this, we focused on 14 grassland species and subjected their seeds to simulated long-term storage by exposing them to artificial ageing conditions (60% rH, 45°C). We then compared plants grown from the aged seeds with plants from fresh seeds in a common garden experiment.
3. Artificially aged seeds germinated later, the developing seedlings had lower survival rates and reduced growth. Adult plants grown from aged seeds flowered later, produced fewer flowers, and had less biomass by the end of the first vegetation period than those from fresh seeds. The effect of the ageing treatment varied between species, but the trend was overall significant across species, with minor differences between perennials and annuals. Interestingly, in perennial plants, the effects vanished or were inverted in the second growing season, with plants growing from aged seeds flowering earlier and producing more biomass.
4. *Synthesis*. Our results show that seed storage affects seedling performance, plant growth, and flowering phenology. These direct storage effects should be considered when using stored seeds for species conservation, ecosystem restoration, or evolutionary research relying on stored seeds.

## INTRODUCTION

Seeds of wild plant species are invaluable reservoirs of biodiversity. As such, they are commonly stored for ex-situ conservation, restoration, or research purposes (Colville, 2017; De Vitis et al., 2020; Etterson et al., 2016). The majority of species have orthodox seeds that tolerate desiccation and freezing (Wyse & Dickie, 2017). Such seeds are optimally stored in conservation seed banks under low temperature and humidity. These conditions preserve seed viability and quality for many decades or even centuries (FAO, 2022; Walters, 2015). However, seeds are also often stored under less optimal conditions, for example, in large quantities before they are used to re-establish diverse plant communities in large-scale restoration projects (Broadhurst et al., 2016; Merritt & Dixon, 2011). The longevity of seeds in storage ultimately depends on how well storage conditions preserve seed quality.

Prolonged storage or exposure to suboptimal environments inevitably causes seed deterioration. Ageing processes, mainly driven by oxidative damage to proteins, lipids, and nucleic acids (El-Maarouf-Bouteau et al., 2011; Fleming et al., 2018; Harman & Mattick, 1976), gradually reduce both seed viability, that is the ability to germinate, and seed vigor, that is the performance of the emerging seedling (Priestley et al., 1985). The rate of seed deterioration varies considerably among and within species (Klepka et al., 2025; Probert et al., 2009), but deterioration is ultimately unavoidable. Most previous studies have focused on the duration of seed viability during storage and predicting seed longevity (Merritt et al., 2014; Satyanti et al., 2018; Walters et al., 2005). There is also evidence that storage delays germination of the surviving seeds (Berjak & Villiers, 1972; Rice & Dyer, 2001; Verma et al., 2003) and causes morphological abnormalities of seedlings such as radicle or cotyledon malformations (Bradford et al., 1993; Delouche & Baskin, 1973). Particularly, cotyledon damage at the seedling stage can impair successful seedling establishment (Hanley & May, 2006).

Emerging evidence suggests that seed deterioration also affects the traits of adult plants grown from surviving seeds. In *Brassica rapa* Fast Plants^®^, plants grown from artificially aged seeds flowered later and had smaller leaves than those from fresh seeds (Franks et al., 2019). Such differences might be caused by two mechanisms: First, storage directly damages the seeds, depletes resources, and consequently creates less favourable conditions for early growth, potentially with lasting effects (Hanley & May, 2006; Waterworth & West, 2023). Second, the seeds that die during storage may not be a random subsample of the original population. If greater tolerance to ageing (i.e., heritable seed longevity) is correlated with other at least partially heritable traits such as seed mass or flowering onset (Larios et al., 2023; Renard et al., 2020), then seed storage can act as an evolutionary filter (Franks et al., 2019). Empirical studies have shown that seed longevity covaries with other seed and life-history traits (Genna et al., 2020; Klepka et al., 2025), suggesting that certain genotypes could be less likely to survive storage, which might result in the “invisible fraction”. This phenomenon refers to the part of the pre-storage population that is no longer observable because the seeds have died. Both processes, the direct storage effects and the invisible fraction, potentially alter traits of both seedlings and adult plants in populations growing from old seeds.

Effects of seed ageing are well documented for germination-associated traits, such as delayed germination, reduced seedling establishment, and morphological abnormalities (Padilha et al., 2022; Rice & Dyer, 2001; Verma et al., 2003). This pattern could be analogous to maternal effects, where conditions experienced by the maternal plant influence offspring traits. Maternal effects typically have the strongest impact during the earliest life stages and often weaken as plants mature (D. A. Roach & Wulff, 1987). By analogy, effects of damage due to ageing may predominantly affect early developmental stages, with their influence weakening over time as individuals recover from the initial stress. Consequently, the effects of seed ageing should be strongest on germination and seedling traits, and weaker on later life-history traits.

It is unknown how widespread the seed deterioration effects are on adult plant traits across species. We are aware of only two studies that have demonstrated the effect of seed ageing (Franks et al., 2019; Ghassemi-Golezani et al., 2010). In *Brassica rapa* Fast Plants^®^, seed ageing delayed flowering and reduced leaf size (Franks et al., 2019). At the same time, the affected traits are adaptive in the wild (Colautti & Barrett, 2013; Giménez-Benavides et al., 2011), and trait modification could affect the performance of the plants from old seeds if used for species reintroductions or ecosystem restoration. Storage effects could also confound results of resurrection studies, because they could mask evolutionary changes in response to recent environmental change (Etterson et al., 2016; Franks et al., 2018). To assess the prevalence and magnitude of these potential problems, we urgently need to understand how seed ageing affects plant traits across a wide range of species.

To fill this gap, we focused on 14 common grassland species from 7 plant families and used artificial ageing to simulate long-term storage and related damage. This approach allows direct comparison between aged and fresh seeds from the same seed lot, and thus, isolates the effect of ageing. We compared germination-related traits between aged and fresh seeds and grew plants from these seeds to compare their growth, flowering phenology, and fitness.

We hypothesise that (1) artificial ageing delays germination, reduces seedling establishment, and results in smaller seedlings, (2) plants growing from aged seeds flower later, produce fewer flowers, lighter seeds, and less biomass at the end of the growing season, and (3) effects of seed ageing decreases over time and be weaker or absent in later stages of plant life.

## MATERIAL AND METHODS

### Study species and seed material

We used seeds of 14 European grassland species representing seven plant families (Table 1). All species have orthodox seeds, which survive desiccation and freezing during storage. The seeds came from European seed producers who produce seeds for restoration. While on-farm propagation may unintentionally select for specific traits (Conrady et al., 2023; Nagel et al., 2019), this effect is likely minor in our samples because most producers do not propagate seeds for many generations (*personal communication*). Farm-propagated seeds further offer the advantage that maternal plants grew under optimal conditions, and the seeds were harvested at full maturity. This reduces variability in seed quality, which massively affects storage behaviour (Sinniah et al., 1998).

**Table 1:** Study species. Durations of the artificial seed ageing are based on preliminary results (Klepka et al., 2025). Viability reduction indicates the relative decrease in the proportion of viable seeds on agar after artificial ageing compared to fresh seeds. Survival reduction indicates the relative decrease in the proportion of healthy seedlings emerging in soil after artificial ageing compared to fresh seeds (combining viability loss and post-germination establishment success).

| Species | Family | Life span | Ageing duration (days) | Viability reduction (%) | Survival reduction (%) | Included in analysis |
| --- | --- | --- | --- | --- | --- | --- |
| *Barbarea vulgaris* | Brassicaceae | Perennial | 4 | 21.82 | 38.36 | Yes |
| *Bromus hordeaceus* | Poaceae | Annual | 4 | 64.04 | - | No<br>(too few seedlings) |
| *Bupleurum rotundifolium* | Apiaceae | Annual | 5 | 23.26 | 33.88 | Yes |
| *Cardamine hirsuta* | Brassicaceae | Annual | 22 | 32.79 | 57.82 | Yes |
| *Centaurea jacea* | Asteraceae | perennial | 18 | 25 | 37.28 | Yes |
| *Dianthus deltoides* | Caryophyllaceae | Perennial | 17 | -1.28 | -3.25 | No<br>(insufficient viability reduction) |
| *Lotus corniculatus* | Fabaceae | Perennial | 25 | 14.58 | 26.83 | No<br>(insufficient viability reduction) |
| *Lychnis flos-cuculi* | Caryophyllaceae | Perennial | 9 | 42.45 | 50 | Yes |
| *Plantago lanceolata* | Plantaginaceae | Perennial | 28 | 43.27 | 43.38 | Yes |
| *Plantago media* | Plantaginaceae | Perennial | 21 | 48.18 | 44.16 | Yes |
| *Senecio vulgaris* | Asteraceae | Annual | 18 | 42.19 | 27.35 | Yes |
| *Stellaria media* | Caryophyllaceae | Annual | 17 | 48.48 | 54.55 | Yes |
| *Thlaspi arvense* | Brassicaceae | Perennial | 8 | 37.07 | 73.33 | No<br>(too few seedlings) |
| *Trifolium campestre* | Fabaceae | Annual | 33 | 5.08 | 43.99 | No<br>(insufficient viability reduction) |

**Table 2:** Results of linear mixed models testing the effect of seed ageing across annual and perennial study species. P-values are based on Type I ANOVA F-tests. Significant effects (p < 0.05) after Benjamini-Hochberg correction are highlighted in bold.

| Year | Life cycle | Model | Estimate | SE | Test Stat | P-value (adjusted) | Effect Size* |
| --- | --- | --- | --- | --- | --- | --- | --- |
| **Germination time** |  |  |  |  |  |  |  |
| 2024 | A | LMM | 3.46 | 0.72 | $F(1,57) = 22.98$ | **< 0.001** | $R^2m = 0.04, R^2c = 0.43$ |
| 2024 | P | LMM | 3.91 | 0.56 | $F(1,41) = 47.99$ | **< 0.001** | $R^2m = 0.05, R^2c = 0.49$ |
| **Seedling establishment** |  |  |  |  |  |  |  |
| 2024 | A | GLMM | -0.67 | 0.24 | $z(1, 46) = -2.74$ | **0.011** | $R^2m = 0.05, R^2c = 0.78$ |
| 2024 | P | GLMM | -0.26 | 0.13 | $z(1, 46) = -2.03$ | 0.063 | $R^2m = 0.08, R^2c = 0.38$ |
| **Seedling biomass** |  |  |  |  |  |  |  |
| 2024 | A | LMM | -0.01 | 0.00 | $F(1,74) = 14.09$ | **0.001** | $R^2m = 0.05, R^2c = 0.73$ |
| 2024 | P | LMM | -0.01 | 0.00 | $F(1,94) = 22.71$ | **< 0.001** | $R^2m = 0.07, R^2c = 0.68$ |
| **Flowering onset** |  |  |  |  |  |  |  |
| 2024 | A | LMM | 1.62 | 1.04 | $F(1,225) = 2.42$ | 0.145 | $R^2m = 0.00, R^2c = 0.90$ |
| 2024 | P | LMM | 3.11 | 0.92 | $F(1,153) = 11.45$ | **0.002** | $R^2m = 0.00, R^2c = 0.97$ |
| 2025 | P | LMM | -5.47 | 1.65 | $F(1,266) = 11.01$ | **0.002** | $R^2m = 0.01, R^2c = 0.87$ |
| **Flower number** |  |  |  |  |  |  |  |
| 2024 | A | LMM | -10.54 | 10.97 | $F(1,229) = 0.92$ | 0.362 | $R^2m = 0.00, R^2c = 0.63$ |
| 2024 | P | LMM | -8.52 | 1.84 | $F(1,150) = 21.36$ | **< 0.001** | $R^2m = 0.10, R^2c = 0.27$ |
| 2025 | P | LMM | 10.46 | 10.98 | $F(1,285) = 0.91$ | 0.362 | $R^2m = 0.00, R^2c = 0.36$ |
| **Seed mass** |  |  |  |  |  |  |  |
| 2024 | A | LMM | -0.06 | 0.04 | $F(1,207) = 2.86$ | 0.119 | $R^2m = 0.00, R^2c = 0.96$ |
| 2024 | P | LMM | -0.07 | 0.04 | $F(1,106) = 2.92$ | 0.119 | $R^2m = 0.00, R^2c = 0.94$ |
| 2025 | P | LMM | 0.01 | 0.05 | $F(1,222) = 0.04$ | 0.850 | $R^2m = 0.00, R^2c = 0.88$ |
| **Adult biomass** |  |  |  |  |  |  |  |
| 2024 | A | LMM | -0.93 | 0.28 | F(1,233) = 11.26 | **0.002** | R<sup>2</sup>m = 0.01, R<sup>2</sup>c = 0.74 |
| 2024 | P | LMM | -4.74 | 0.73 | F(1,293) = 42.62 | **< 0.001** | R<sup>2</sup>m = 0.09, R<sup>2</sup>c = 0.38 |
| 2025 | P | LMM | 2.48 | 1.07 | F(1,286) = 5.35 | **0.035** | R<sup>2</sup>m = 0.01, R<sup>2</sup>c = 0.29 |

Note: Models: LMM = Linear Mixed Model, GLMM = Generalized Linear Mixed Model
\* R<sup>2</sup>m = marginal R<sup>2</sup> (fixed effects only), R<sup>2</sup>c = conditional R<sup>2</sup> (fixed + random effects)

### Artificial ageing treatment

We applied artificial seed ageing to compare individuals from the same seed lot that differed only in their simulated age. Artificial ageing is a standard method to study seed longevity (e.g. Hay et al., 2019; Merritt et al., 2014; Probert et al., 2009; Tausch et al., 2019). In our study, this approach ensured that any differences between plants grown from aged and fresh seeds reflected the impact of the ageing treatment rather than different evolutionary pasts. We exposed the seeds to 45°C and 60% relative humidity in a climate-controlled cabinet (Rumed ®, Rubarth Apparate GmbH, Laatzen, Germany) to simulate storage. These conditions accelerate chemical processes typical of natural ageing during long-term storage and enhance seed deterioration (Delouche & Baskin, 1973; Rajjou & Debeaujon, 2008). Although artificial ageing does not fully replicate seed ageing under natural conditions, both impose stress on seeds, leading to a viability decline (Silva et al., 2023). For the purpose of our study, artificial seed ageing represents stress experienced in individuals in the seed stage, which is similar to stress experienced during storage. It can therefore be used to investigate how storage-related stress affects the performance of plants emerging from aged seeds.

In a previous experiment (Klepka et al., 2025), we estimated seed deterioration curves of the seed lots and used them to determine the seed lot-specific time under artificial ageing conditions until seed viability was reduced by 50%. We then exposed the seeds to artificial ageing for the specified time with the aim of reducing viability to 50% of the viability of the original seed lot (Table 1). Before ageing, all seeds were kept on a laboratory bench in paper bags for two months to equilibrate with surrounding humidity and temperature (ca 21°C, 50% rH).

### Germination trial

We placed the seeds in batches of 16 per replicate in Petri dishes. For each species, we had 4 replicates for fresh seeds (4 x 16 = 64 seeds) and 8 replicates for aged seeds (8 x 16 = 128 seeds), in total 2,688 seeds (14 species x (64 + 128)). The Petri dishes contained 1% agar with an anti-mould agent (0.0835% Difenoconazol) and gibberellic acid (0.00025%) to break dormancy. We then incubated the dishes in a climate-controlled cabinet (Rumed®, Rubarth Apparate GmbH, Laatzen, Germany) in a fully randomized design and under a 14h light / 10h dark cycle at 20°C / 10°C. We scored signs of viability on the level of individual seeds three times per week. When a seed had formed either a radicle (≥ 2mm, Millennium Seed Bank, 2022) or cotyledons, we considered it viable. We removed mouldy seeds immediately to prevent further contamination and considered it not viable.

### Common garden experiment

We grew adult plants from fresh and artificially aged seeds and compared them in a common environment. For each species, we sowed 25 fresh seeds and 50 aged seeds into germination trays in 4 and 6 replicates, for a total of 100 and 300 seeds, respectively. We used a larger number of aged seeds per tray and a higher number of replicates to compensate for lower seed viability and because we expected higher seedling mortality for the aged seeds (Padilha et al., 2022). The seeding trays were filled with standard seeding substrate (Einheitserde^®^ ED73). We placed the trays in an unheated greenhouse for germination in a randomized design and counted established seedlings. Once the seedlings developed their first true leaves, we randomly selected 60 individuals per treatment and species and transferred them to QuickPots^®^. The remaining seedlings were kept in trays, and when most of them reached a size of 10 cm, we harvested 10 replicates of 2 to 3 seedlings across all trays, dried them at 70°C, and weighed them to determine seedling biomass.

Because seedling size varied among species, we adjusted the number of seedlings per replicate accordingly: For species with larger seedlings, we harvested 10 replicates of 2 random seedlings each; for species with smaller seedlings, we harvested 10 replicates of 3 random seedlings each. We then calculated the average seedling weight per replicate. Seedlings of each species from both aged and fresh seeds were transplanted and later harvested on the same day to avoid the confounding effect of transplanting or harvesting.

Once the plants in the QuickPots^®^ had reached a height of at least 10cm, we selected 30 random individuals per treatment and species and transplanted them into 2.5L pots filled with standard potting soil. We arranged the plants randomly in a common garden and watered them as needed. The common garden was located in a net house that excluded pollinators. We recorded flowering phenology as the day of the first open flower per individual. We manually cross-pollinated the flowers within a treatment to obtain seeds. At the end of the growing season, we counted all inflorescences that each plant produced during the season, collected ripe seeds, harvested aboveground biomass, dried it at 70°C, and weighed it. We determined seed weight by weighing 3 replicates of 20 seeds for the maternal plant, and we report it as thousand-seed weight.

Annual species included in the experiment completed their life cycle in the autumn of the first growing season. Perennial species regrew after harvest of aboveground biomass. We maintained them in the common garden and recorded the same adult traits, also in the second growing season. This allowed us to test for the effects of seed ageing in later plant life, and obtain the onset of flowering, number of flowers, and seed weight for all perennial species, because not all species started to flower in the first season. At the beginning of the second growing season, we fertilized the pots with slow-release fertilizer (Osmocote^®^). We recorded all data using the GridScore NEXT app (Raubach et al., 2022).

### Focal traits

We focused on the effect of seed ageing on germination and adult traits. Germination traits included germination timing, measured as time until signs of viability for each seed, and seedling establishment, measured as the number of viable seeds that successfully established as seedlings. For the latter, the number of viable seeds per species and treatment was estimated from the germination trial data, allowing us to calculate how many viable seeds per seeding tray failed to establish.

Adult traits were all measured at the individual level and included the onset of flowering, defined as the day of the year when the first flower showed visible stamens, the number of flowers or inflorescences, the total aboveground biomass at the end of the growing season, and seed weight, measured as the thousand-seed weight.

For adult traits in perennials, we collected data from two consecutive growing seasons. We recorded biomass for all species in both years. However, for the other adult traits (flowering onset, flower number, and seed mass), we only obtained data for some species. Specifically, not all individuals of perennials had started to flower in the first growing season or produced a sufficient number of seeds. In the first growing season, we thus had flowering time and number of inflorescences for only part of the perennial species. To have reliable data, we included first-year data for flowering traits only if at least 20% of individuals per treatment had flowered. In the second growing season, we recorded all the traits for all perennials. Due to this unbalanced data structure, we treated traits from the two growing seasons as independent in all analyses.

### Statistical analyses

First, we tested whether the artificial ageing treatment indeed reduced seed viability to the aimed 50% of the fresh seeds. For this, we calculated seed viability reduction caused by ageing as the relative decrease in viability in aged seeds compared to fresh seeds, based on germination trials in Petri dishes. Contrary to our aim of 50%, the viability reduction varied among seed lots between 0 and 64% (Table 1). To ensure that seed ageing indeed had a substantial impact on the seed quality, we excluded species where viability declined by less than 20% from further analyses (*Dianthus deltoides*, *Lotus corniculatus*, *Trifolium campestre*).

Second, we analysed how the ageing treatment affected germination and adult plant traits of each species separately. For each species, trait, and growing season (the latter in perennials only), we fitted an individual model with treatment (artificial ageing vs. control) as explanatory variable. For germination time, we fitted linear mixed models with the days until viability signs for each seed as the response and the Petri dish number as random effect. For seedling establishment, we fitted generalised linear models with binomial error distributions and probit-link functions. As response variable, we used the number of successfully established vs. viable but non-established seedlings (estimated from the germination trial), specified with the cbind() command. For seedling biomass, we fitted linear models with individual seedling biomass as the response. For adult plant traits, we fitted linear models with the respective traits as response. To control for the false discovery rate caused by multiple testing, we adjusted the resulting p-values using the Benjamini-Hochberg procedure (Benjamini & Hochberg, 1995).

Third, we tested the overall effects of seed ageing across all study species. We considered annuals and perennials separately due to differences in their life cycle and data structures. For perennials, we analysed adult traits from each growing season separately because of many missing values in the first season. We thus fitted linear mixed models for each life cycle group (annual and perennial) and, for perennials, each growing season. The response variable was the trait of interest, with the ageing treatment as the fixed explanatory variable, and species identity as the random factors. For germination time and seedling establishment, we also added Petri dish and seeding tray as random factor, respectively. The error structures were the same as those in the species-specific models for each trait.

We used the R-packages *lme4* (Bates et al., 2015), *emmeans* (Piaskowski, 2025), *lmerTest* (Kuznetsova et al., 2017), and *car* (Fox et al., 2024) to fit linear models and obtain model parameters. We checked model assumptions using the *performance* package (Lüdecke et al., 2021) and found them to be reasonably met. All data analyses were performed using R version 4.4.3 (R Development Core Team, 2024).

## RESULTS

Germination traits were consistently affected by seed ageing. Across all species, artificially aged seeds germinated later, were less likely to establish into healthy seedlings (significant for perennials only), and produced seedlings with less biomass. At the species level, 6 of 11 species germinated significantly later from aged seeds, 4 out of 10 species showed reduced seedling establishment and/or produced smaller seedlings. A notable exception was *Senecio vulgaris*, in which seedlings from aged seeds established better than seedlings from fresh seeds.

Artificial ageing also affected adult traits, although this effect was less frequent. Across species, perennials growing from artificially aged seeds flowered later, produced fewer flowers, and had less biomass at the end of the first growing season. There was no overall effect of seed ageing on the seed weight of the offspring. In annuals, the seed ageing treatment only significantly reduced the final above-ground biomass (Figure 1 and S1). At the species level, the effect of seed ageing on adult traits varied, with some species being affected in one or two traits. In 4 species, 1 annual and 3 perennials, seed ageing did not significantly affect any of the adult traits. Notably, the effect was most consistent for both *Plantago* species, which produced both fewer flowers and less biomass when plants were grown from aged seeds. The most affected species was *Plantago media*, which was affected in all measured adult traits (Figure 1).

**Figure 1:**
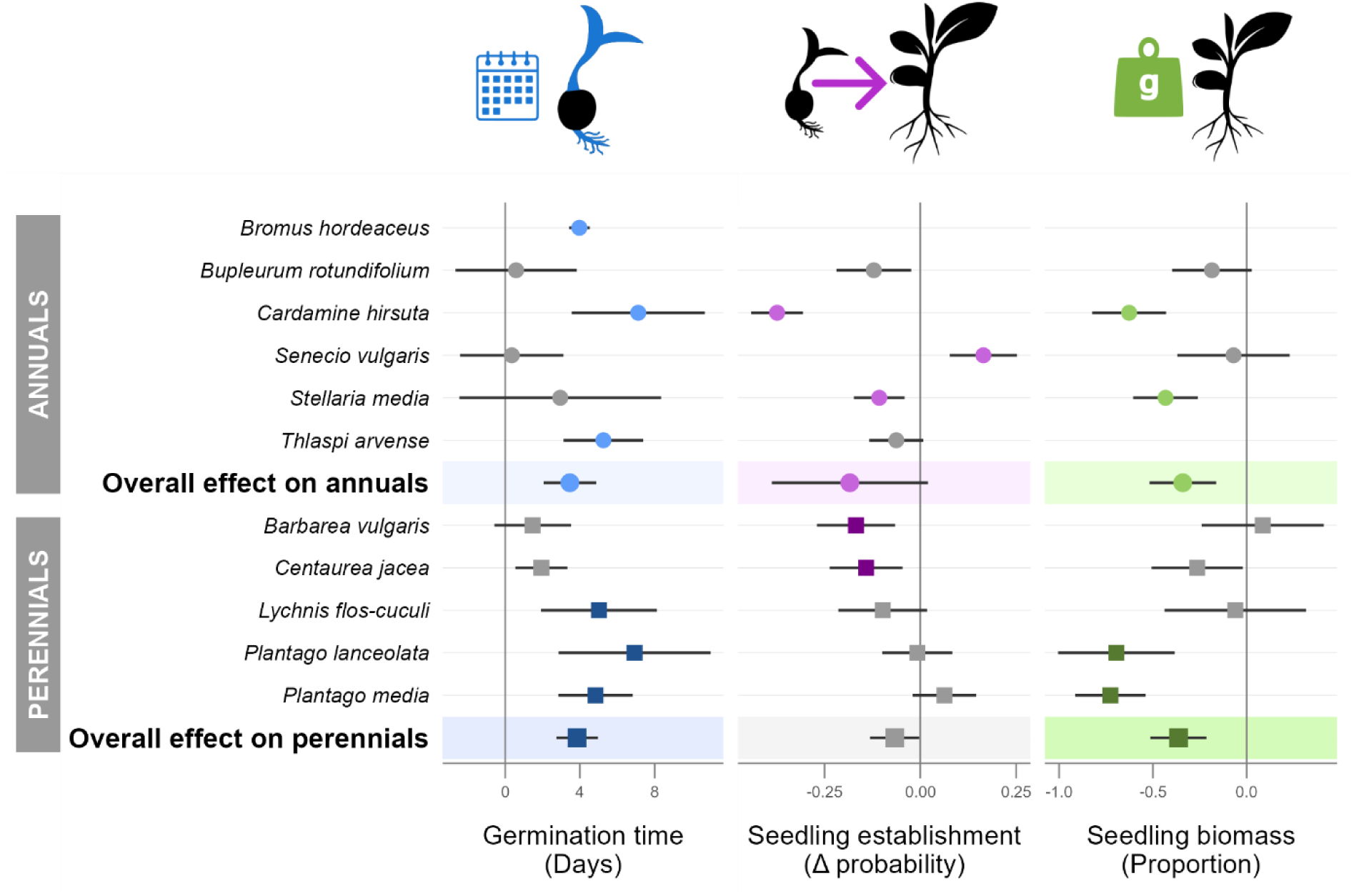
Effect of seed ageing on germination-related traits at the species level, shown separately for annuals and perennials. Points show effect sizes, error bars 95% confidence intervals. For germination time, the effect size is the model estimate of the ageing treatment (in days). For seedling establishment, it is the absolute difference in establishment probability between aged and control seeds (back-transformed from probit scale). For seedling biomass, it is the proportional change relative to the control, calculated as the treatment estimate divided by the intercept. Positive values indicate increases (or delays) in response to the ageing treatment. Coloured points and background shadings indicate significance (p < 0.05). Species are grouped by life cycle: Annuals (upper section, circles, lighter colour) and perennials (lower section, squares, darker colour). See Tables 2 and S1 for individual model results.

**Figure 2:**
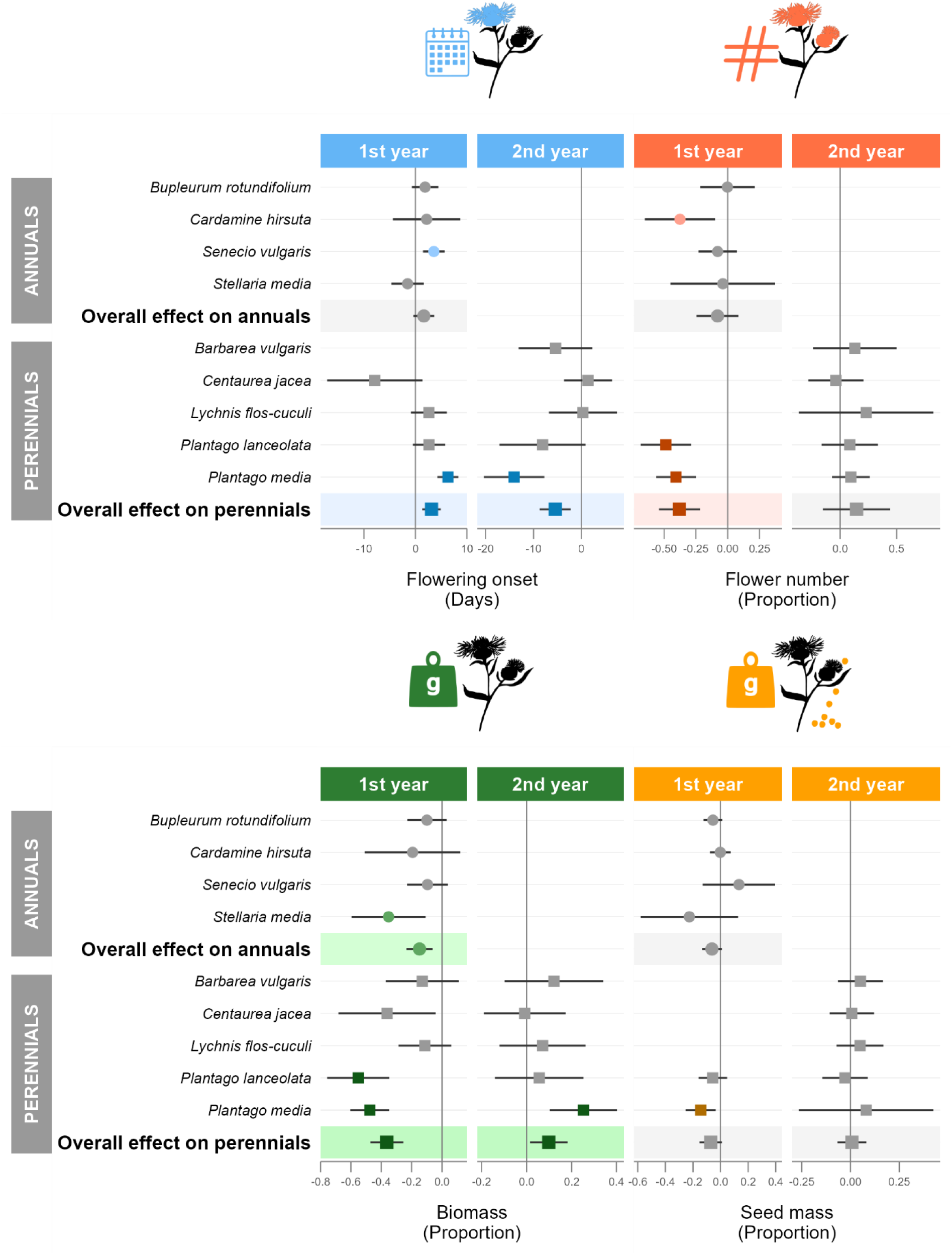
Effect of seed ageing on adult plant traits at the species level, shown separately for annuals and perennials. Points show effect sizes, error bars 95% confidence intervals. For flowering onset, the effect size is the model estimate of the ageing treatment (in days). For the flower number, the final biomass and the seed mass, it is the proportional change relative to the control, calculated as the treatment estimate divided by the intercept. Positive values indicate increases (or delays) in response to the ageing treatment. Coloured points and background shadings indicate significance (p < 0.05). Species are grouped by life cycle: Annuals (upper section, circles, lighter colour) and perennials (lower section, squares, darker colour). See Tables 2 and S1 for individual model results.

Seed ageing partially affected adult traits of perennials in the second growing season, but the effects were weaker or in the opposite direction than in the first growing season. Across all perennials, plants from aged seeds flowered earlier and had more biomass than the control plants. The number of flowers and the seed mass did not differ significantly (Figure 1 and S2). At the species level, *Plantago media* was the only species in which adult plants grown from aged seeds differed significantly from those grown from fresh seeds, specifically, they flowered earlier and had a higher aboveground dry biomass (Figure 1).

## Discussion

Seeds stored in conservation seed banks and other facilities inevitably deteriorate and lose both viability and vigour. Here, we show that even when seeds survive storage, seedlings from aged seeds perform worse during early development than those from fresh seeds, and the effects of ageing can persist into the adult stage. The magnitude and significance of the ageing effects varied between species and growth forms, yet the direction mostly pointed towards reduced performance of plants from aged seeds. Our results demonstrate that seed ageing causes direct and lasting carry-over effects on plants growing from the aged seeds, with consequences extending far beyond germination.

Seed ageing most consistently affected germination traits. Aged seeds germinated later, the seedlings established less successfully, and were smaller than those from fresh seeds. This aligns with our first hypothesis and is most likely because seeds must repair accumulated biochemical damage before radicle emergence. Delayed germination of aged seeds aligns with previous studies (Berjak & Villiers, 1972; Padilha et al., 2022) and is likely due to the time required to repair ageing-induced damage before germination can proceed (Berjak & Villiers, 1972). Oxidative processes during ageing damage nucleic acids, lipids, and proteins (Padilha et al., 2022; Rajjou et al., 2008), reduce genome integrity and the ability to mobilise reserves (Rajjou et al., 2008). This ageing-induced cellular damage, together with resource depletion and higher proportions of morphologically abnormal seedlings after ageing (Clerkx et al., 2003; Rehmani et al., 2023; Tesnier et al., 2002), likely contributed to significant reductions in seedling establishment. Our results demonstrate that viability loss during ageing does not fully capture the eventual loss of individuals from the population of germinated seeds, because more individuals will die during the seedling stage. Standard germination trials may therefore systematically overestimate the effective viability of collections of stored seeds.

Seed ageing affected adult plant traits. Generally, plants from aged seeds flowered later, produced fewer flowers, and less biomass at the end of the first growing season. These results are consistent with findings in *Brassica rapa* (Franks et al., 2019) and *Brassica napus* (Ghassemi-Golezani et al., 2010). Underlying mechanisms include disruption of endogenous cues and molecular pathways regulating flowering time (Amasino & Michaels, 2010; Poethig & Fouracre, 2024). Additionally, ageing-induced delays in germination and reductions in seedling size may have delayed reaching the size or developmental thresholds necessary for flowering onset (Poethig & Fouracre, 2024). Thus, reduced flower number and the lower aboveground dry biomass at the end of the first growing season possibly reflect the cumulative impact of all earlier disadvantages caused by the ageing treatment.

The ageing affected adult plant traits also in the second growing season in some but not all perennial plants. We expected that the effect of ageing would be weaker or absent in later life stages, which was true only for some species. However, in the species where the effect of ageing persisted, it was opposite to the first growing season. For example, perennials from aged seeds flowered later than plants from fresh seeds in the first growing season but earlier in the second growing season. A possible mechanism behind this observation could be a shift in resource allocation: The early-life constraints of delayed germination and reduced growth, driven by the need for repair and resource depletion, may have triggered higher investment in below-ground organs during the first growing season (Moriuchi & Winn, 2005), which could represent an advantage in the second season. Such a pattern might also be the result of overcompensatory responses to early stress, where individuals who experience restrictions perform better later in life compared to unstressed individuals (Roitberg et al., 2024). This finding suggests that extrapolation from short-term effects to long-term effects may be misleading.

The response of traits to seed ageing differed substantially among species. The effect of seed ageing on germination traits was the most consistent one, and the majority of species showed some signs of reduced performance. One notable exception was *Senecio vulgaris,* which showed a better seedling establishment after ageing. This could result from selective filtering during the ageing treatment, where weaker individuals die first, and remaining seeds represent a subset of particularly fit individuals with a higher survival probability. In adult traits, the overall effects pointed in the same direction for annuals and perennials, but the effects were more frequently significant for perennials. This can be an effect of power, because our experiments included 5 perennials and only 4 annuals. Additionally, the effect on perennials was driven by the two *Plantago* species, and this genus may be particularly sensitive to seed ageing. Nevertheless, our results demonstrate that the strength of ageing effects on subsequent life-history traits is species-specific. Identifying the drivers behind the species differences would require experiments with many more species across diverse families (van Kleunen et al., 2014).

The ageing effects on plant traits are possibly not only driven by the direct chemical changes in the seed during ageing, but also by non-random seed mortality. Survival probability during storage may be genetically correlated with other traits such as seed size, flowering phenology or dormancy (Franks et al., 2019; Nguyen et al., 2012; Renard et al., 2020). Non-random loss of seeds would thus cause a shift in the trait means of the correlated traits. This phenomenon is called the “invisible fraction”, and it refers to trait values that cannot be measured because the individuals died during the seed storage (Franks et al., 2019; Weis, 2018). While reduced plant performance in the first growing season could principally be the result of the invisible fraction (Franks et al., 2019), the reversal of negative effects in perennials in the second season is more consistent with direct physiological damage than with loss of genotype. Nevertheless, to completely disentangle the invisible fraction from direct storage-induced effects will require growing a second generation free of direct storage effects.

Our study accelerated seed ageing by exposing seeds to high temperature and humidity. This method is widely accepted for studying seed longevity, particularly for comparative studies and monitoring (Hay et al., 2019, 2022). Exposing seeds to high temperatures and humidity is assumed to accelerate similar chemical processes in seeds to those typical of natural ageing during long-term storage (Delouche & Baskin, 1973; Rajjou et al., 2008). However, artificial ageing does not fully mirror natural ageing (Gianella et al., 2022; Hay et al., 2019; T. Roach et al., 2018). Despite these limitations, both natural and artificial ageing impose stress on seeds and consistently reduce seed viability and vigour (Silva et al., 2023). Although the underlying processes are not identical, artificial ageing represents the most practical and widely used approach for studying seed deterioration (Hay et al., 2019; Merritt et al., 2014; Probert et al., 2009; Tausch et al., 2019). It can therefore be used to investigate how storage-related stress affects the performance of plants emerging from aged seeds. However, caution is required when precisely inferring the consequences of long-term storage for different species or genotypes.

The hot and humid conditions during the artificial ageing treatment could have caused an acute heat and moisture stress. Such early-life stresses can influence later plant performance and their effects sometimes persist for months, or even across generations, likely through epigenetic modifications (Crisp et al., 2016; Lodhi et al., 2025; Zi et al., 2024). However, such a stress memory may be relatively rare (Crisp et al., 2016). Additionally, there is evidence showing that seedlings from naturally aged seeds perform worse than seedlings from fresh seeds (Rice & Dyer, 2001; Verma et al., 2003), which aligns with our results. It is thus unlikely that our findings are an artifact of the artificial seed ageing method.

### Implications for practice

Our findings have important implications for any application that uses stored seeds. In conservation seed banks, it would be beneficial to supplement regular germination tests for viability monitoring with seedling establishment assays, as viability tests alone may overestimate the number of adult plants that can be obtained from the seeds. This would give a more accurate proxy for seed survival.

When using stored seed for reintroductions of rare species or for ecosystem restoration, we recommend using seeds as fresh as possible or employing a refresher generation. Not only because old seeds have reduced germination and lower seedling establishment rates, but also because plants from old seeds may have altered adult traits like reduced growth or later flowering. These traits are often adaptive in the wild (Colautti & Barrett, 2013; Giménez-Benavides et al., 2011), and their alteration could impact reintroduction or restoration success. When using stored seeds is unavoidable, practitioners should anticipate reduced establishment rates and increase seed density accordingly.

For resurrection studies that rely on stored seeds to detect evolutionary responses to environmental change, refresher generations remain essential before comparing ancestors with descendants, as it is already standard practice to remove direct physiological effects (Franks et al., 2018, 2019). However, we acknowledge that this is not always feasible for all experimental settings or species, and several resurrection studies therefore directly compare ancestral seeds with their present-day descendants without refresher generation (e.g., White et al., 2023). In such cases, differences may partly reflect direct storage effects rather than evolutionary change. Furthermore, if storage acts as a selective filter (invisible fraction), even refresher generations may not fully account for storage effects. We recommend using seed lots with high germination and establishment rates to minimize loss of genotypes and cautious interpretation of results from resurrection studies.

## Author contribution

AB conceived the ideas and designed the methodology; LK, SK, EH, and PC collected the data; LK analysed the data; AB and LK led the writing of the manuscript. All authors contributed critically to the drafts and gave final approval for publication.

## Acknowledgement

We thank Bendrik Baumeister, Emelie Hähner, and Christina Mengel for technical support, and the Marburg University Research Academy (MARA) for providing a scholarship that supported the research.

## Conflict of interest statement

The authors declare no conflicts of interest.

## Data accessibility statement

The data supporting the findings are not yet publicly available at the time of submission, but will be deposited in an appropriate public repository upon acceptance of the manuscript. Data will be made available to editors and reviewers upon reasonable request.

---

**Figure S1:**
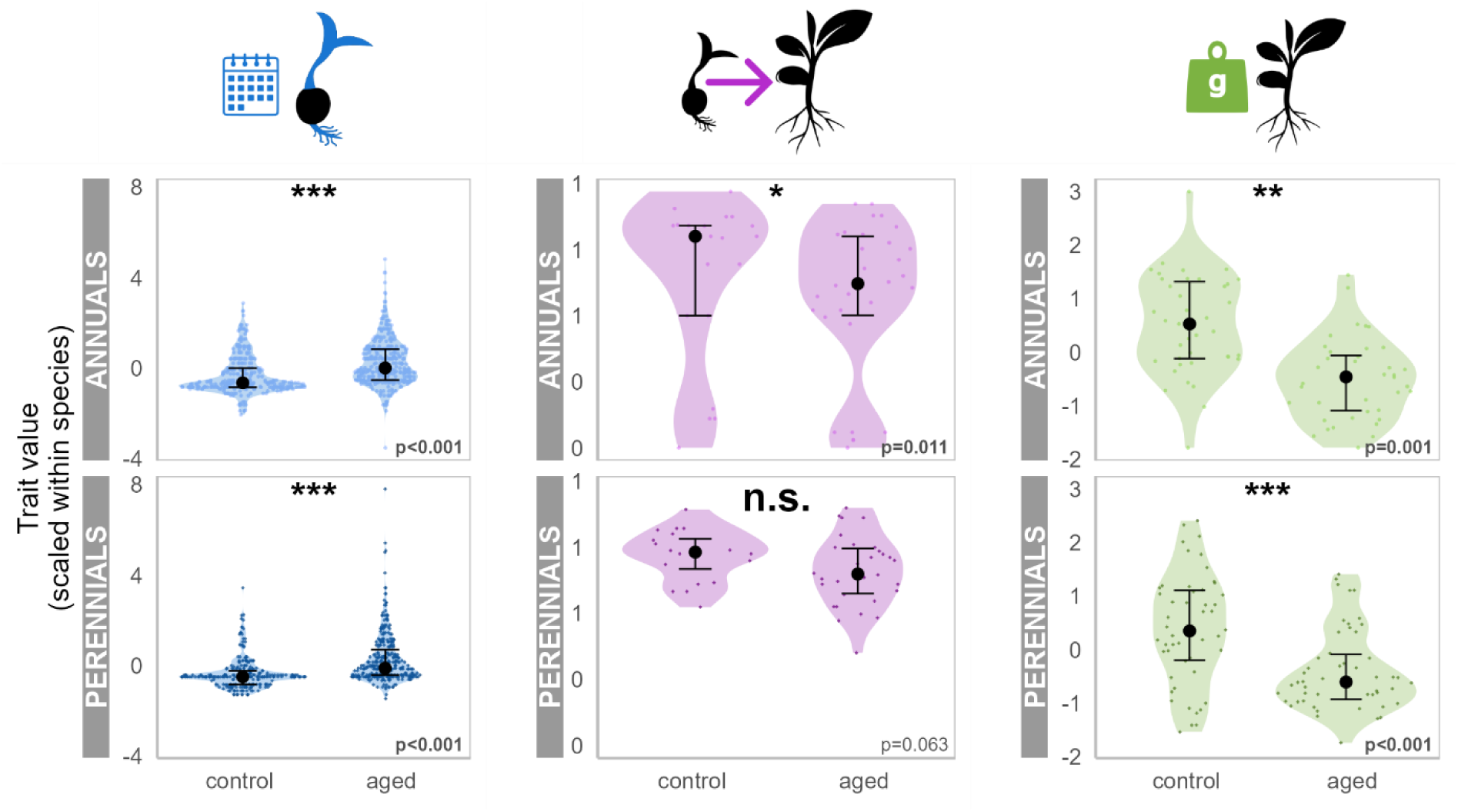
Effect of seed ageing on germination traits across all species separated by life cycle. Each point is an individual with its value scaled within the species. Black dots are median values with 25^th^-75^th^ percentiles.

**Figure S2:**
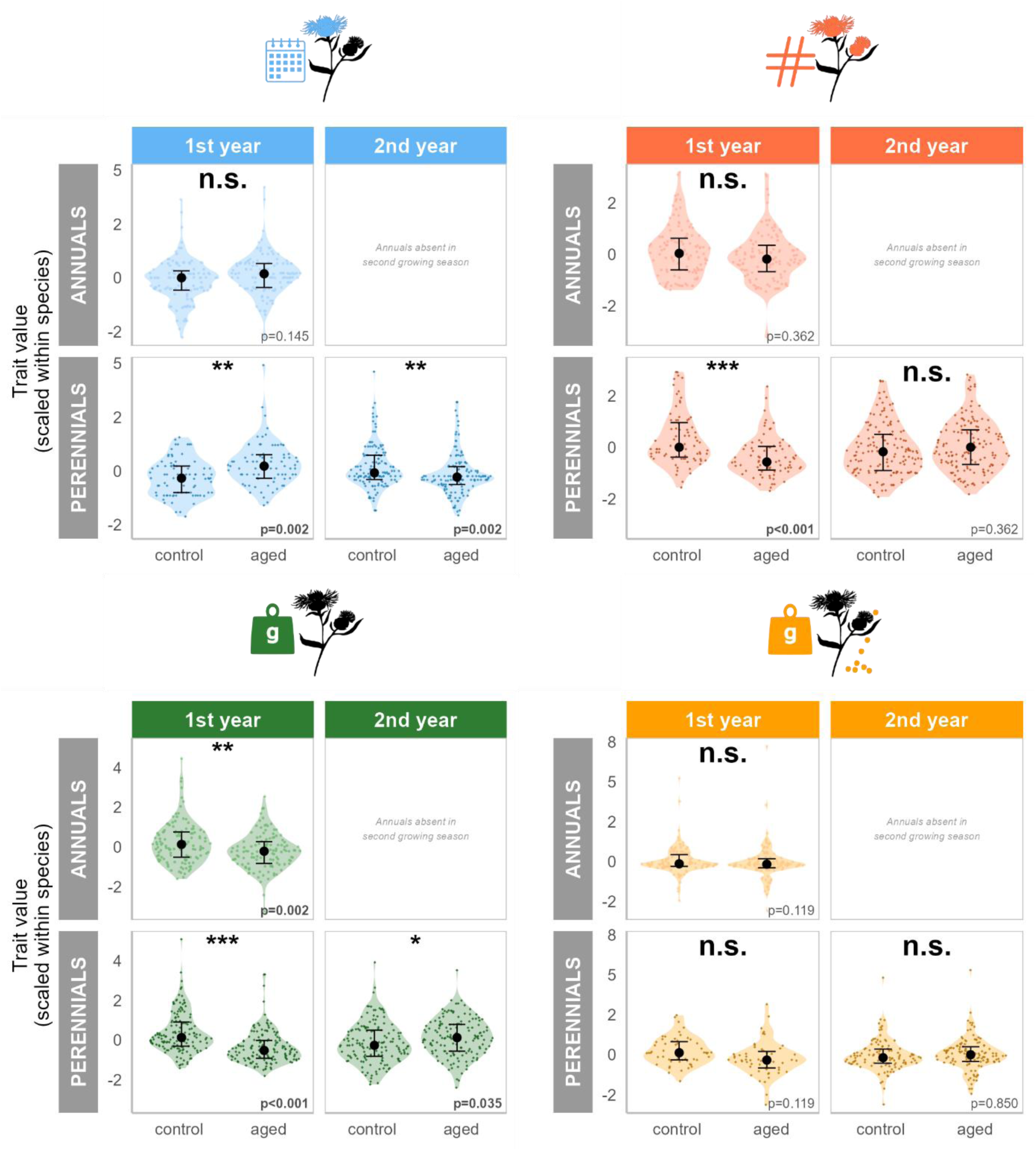
Effect of seed ageing on adult plant traits across all species separated by life cycle and growing season. Each point is an individual with its value scaled within the species. Black dots are median values with 25^th^-75^th^ percentiles.

**Table S1:** Results of linear models testing the effect of seed ageing on germination and adult plant traits for each species. P-values are based on Type I ANOVA F-tests. Significant effects (p < 0.05) after Benjamini-Hochberg correction are highlighted in bold.

| Species | Year | Model | Estimate | SE | Test Stat | P-value<br>(adjusted) | Effect Size* |
| --- | --- | --- | --- | --- | --- | --- | --- |
| **Germination time** |  |  |  |  |  |  |  |
| *Barbarea vulgaris* | 2024 | LMM | 1.42 | 1.04 | $F(1, 9) = 1.86$ | 0.366 | $R^2m = 0.03, R^2c = 0.17$ |
| *Bromus hordeaceus* | 2024 | LMM | 3.97 | 0.29 | $F(1, 12) = 192.63$ | **< 0.001** | $R^2m = 0.68, R^2c = 0.68$ |
| *Bupleurum rotundifolium* | 2024 | LMM | 0.58 | 1.66 | $F(1, 9) = 0.12$ | 0.839 | $R^2m = 0.00, R^2c = 0.22$ |
| *Cardamine hirsuta* | 2024 | LMM | 7.16 | 1.85 | $F(1, 9) = 14.99$ | **0.015** | $R^2m = 0.15, R^2c = 0.21$ |
| *Centaurea jacea* | 2024 | LMM | 1.85 | 0.69 | $F(1, 8) = 7.12$ | 0.082 | $R^2m = 0.09, R^2c = 0.16$ |
| *Lychnis flos-cuculi* | 2024 | LMM | 4.83 | 1.52 | $F(1, 8) = 10.14$ | **0.039** | $R^2m = 0.15, R^2c = 0.24$ |
| *Plantago lanceolata* | 2024 | LMM | 7.22 | 2.06 | $F(1, 108) = 12.43$ | **0.003** | $R^2m = 0.10$ |
| *Plantago media* | 2024 | LMM | 5.16 | 1.03 | $F(1, 109) = 25.36$ | **< 0.001** | $R^2m = 0.19$ |
| *Senecio vulgaris* | 2024 | LMM | 0.11 | 1.37 | $F(1, 9) = 0.01$ | 0.964 | $R^2m = 0.00, R^2c = 0.02$ |
| *Stellaria media* | 2024 | LMM | 2.94 | 2.81 | $F(1, 64) = 1.14$ | 0.454 | $R^2m = 0.02$ |
| *Thlaspi arvense* | 2024 | LMM | 5.42 | 1.09 | $F(1, 128) = 24.74$ | **< 0.001** | $R^2m = 0.16$ |
| **Seedling establishment** |  |  |  |  |  |  |  |
| *Barbarea vulgaris* | 2024 | GLM | -0.17 | 0.05 | $z(8) = -2.94$ | **0.011** | McFadden $R^2$ (adj)= 0.97 |
| *Bupleurum rotundifolium* | 2024 | GLM | -0.12 | 0.05 | $z(8) = -2.17$ | 0.076 | McFadden $R^2$ (adj)= 0.96 |
| *Cardamine hirsuta* | 2024 | GLM | -0.37 | 0.03 | $z(8) = -0.01$ | **< 0.001** | McFadden $R^2$ (adj)= 0.99 |
| *Centaurea jacea* | 2024 | GLM | -0.14 | 0.05 | $z(8) = -2.64$ | **0.024** | McFadden $R^2$ (adj)= 0.97 |
| *Lychnis flos-cuculi* | 2024 | GLM | -0.10 | 0.06 | $z(8) = -1.61$ | 0.249 | McFadden $R^2$ (adj)= 0.96 |
| *Plantago lanceolata* | 2024 | GLM | -0.01 | 0.05 | $z(8) = -0.17$ | 0.935 | McFadden $R^2$ (adj)= 0.95 |
| *Plantago media* | 2024 | GLM | 0.06 | 0.04 | $z(8) = 1.55$ | 0.262 | McFadden $R^2$<br>(adj)= 0.95 |
| *Senecio vulgaris* | 2024 | GLM | 0.16 | 0.04 | $z(8) = 3.97$ | **< 0.001** | McFadden $R^2$<br>(adj)= 0.97 |
| *Stellaria media* | 2024 | GLM | -0.11 | 0.03 | $z(8) = -0.01$ | **0.009** | McFadden $R^2$<br>(adj)= 0.94 |
| *Thlaspi arvense* | 2024 | GLM | -0.06 | 0.04 | $z(8) = -1.89$ | 0.156 | McFadden $R^2$<br>(adj)= 0.96 |
| **Seedling biomass** |  |  |  |  |  |  |  |
| *Barbarea vulgaris* | 2024 | LM | 0.00 | 0.00 | $F(1,18) = 0.27$ | 0.752 | $R^2 = 0.01$ (adj. - 0.04) |
| *Bupleurum rotundifolium* | 2024 | LM | -0.00 | 0.00 | $F(1,17) = 2.94$ | 0.249 | $R^2 = 0.15$ (adj. 0.10) |
| *Cardamine hirsuta* | 2024 | LM | -0.03 | 0.00 | $F(1,18) = 38.83$ | **< 0.001** | $R^2 = 0.68$ (adj. 0.67) |
| *Centaurea jacea* | 2024 | LM | -0.01 | 0.00 | $F(1,18) = 4.54$ | 0.127 | $R^2 = 0.20$ (adj. 0.16) |
| *Lychnis flos-cuculi* | 2024 | LM | -0.00 | 0.00 | $F(1,18) = 0.10$ | 0.840 | $R^2 = 0.01$ (adj. - 0.05) |
| *Plantago lanceolata* | 2024 | LM | -0.02 | 0.00 | $F(1,18) = 19.21$ | **0.002** | $R^2 = 0.52$ (adj. 0.49) |
| *Plantago media* | 2024 | LM | -0.01 | 0.00 | $F(1,18) = 57.91$ | **< 0.001** | $R^2 = 0.76$ (adj. 0.75) |
| *Senecio vulgaris* | 2024 | LM | -0.00 | 0.01 | $F(1,18) = 0.21$ | 0.764 | $R^2 = 0.01$ (adj. - 0.04) |
| *Stellaria media* | 2024 | LM | -0.00 | 0.00 | $F(1,18) = 24.17$ | **< 0.001** | $R^2 = 0.57$ (adj. 0.55) |
| **Flowering onset** |  |  |  |  |  |  |  |
| *Bupleurum rotundifolium* | 2024 | LM | 1.86 | 1.31 | $F(1,56) = 2.01$ | 0.311 | $R^2 = 0.03$ (adj. 0.02) |
| *Cardamine hirsuta* | 2024 | LM | 2.17 | 3.34 | $F(1,58) = 0.42$ | 0.668 | $R^2 = 0.01$ (adj. - 0.01) |
| *Centaurea jacea* | 2024 | LM | -7.90 | 4.73 | $F(1,13) = 2.78$ | 0.262 | $R^2 = 0.18$ (adj. 0.11) |
| *Lychnis flos-cuculi* | 2024 | LM | 2.60 | 1.78 | $F(1,22) = 2.14$ | 0.311 | $R^2 = 0.09$ (adj. 0.05) |
| *Plantago lanceolata* | 2024 | LM | 2.62 | 1.60 | $F(1,57) = 2.67$ | 0.249 | $R^2 = 0.04$ (adj. 0.03) |
| *Plantago media* | 2024 | LM | 6.30 | 1.04 | $F(1,58) = 36.41$ | **< 0.001** | $R^2 = 0.39$ (adj. 0.38) |
| *Senecio vulgaris* | 2024 | LM | 3.57 | 1.07 | $F(1,58) = 11.04$ | **0.007** | $R^2 = 0.16$ (adj. 0.15) |
| *Stellaria media* | 2024 | LM | -1.54 | 1.61 | $F(1,50) = 0.91$ | 0.499 | $R^2 = 0.02$ (adj. - 0.00) |
| *Barbarea vulgaris* | 2025 | LM | -5.40 | 3.93 | F(1,51) =<br>1.88 | 0.324 | R<sup>2</sup> = 0.04 (adj.<br>0.02) |
| *Centaurea jacea* | 2025 | LM | 1.38 | 2.57 | F(1,59) =<br>0.29 | 0.740 | R<sup>2</sup> = 0.00 (adj. -<br>0.01) |
| *Lychnis flos-cuculi* | 2025 | LM | 0.33 | 3.63 | F(1,40) =<br>0.01 | 0.964 | R<sup>2</sup> = 0.00 (adj. -<br>0.02) |
| *Plantago lanceolata* | 2025 | LM | -8.11 | 4.59 | F(1,54) =<br>3.13 | 0.209 | R<sup>2</sup> = 0.05 (adj.<br>0.04) |
| *Plantago media* | 2025 | LM | -14.07 | 3.22 | F(1,58) =<br>19.14 | **< 0.001** | R<sup>2</sup> = 0.25 (adj.<br>0.24) |
| **Flower number** |  |  |  |  |  |  |  |
| *Bupleurum rotundifolium* | 2024 | LM | -0.10 | 4.39 | F(1,56) =<br>0.00 | 0.981 | R<sup>2</sup> = 0.00 (adj. -<br>0.02) |
| *Cardamine hirsuta* | 2024 | LM | -13.79 | 5.17 | F(1,56) =<br>7.10 | **0.034** | R<sup>2</sup> = 0.11 (adj.<br>0.10) |
| *Centaurea jacea* | 2024 | LM | 3.10 | 8.63 | F(1,13) =<br>0.13 | 0.839 | R<sup>2</sup> = 0.01 (adj. -<br>0.07) |
| *Lychnis flos-cuculi* | 2024 | LM | 5.77 | 3.74 | F(1,21) =<br>2.37 | 0.281 | R<sup>2</sup> = 0.10 (adj.<br>0.06) |
| *Plantago lanceolata* | 2024 | LM | -16.54 | 3.43 | F(1,56) =<br>23.23 | **< 0.001** | R<sup>2</sup> = 0.29 (adj.<br>0.28) |
| *Plantago media* | 2024 | LM | -8.90 | 1.74 | F(1,57) =<br>26.06 | **< 0.001** | R<sup>2</sup> = 0.31 (adj.<br>0.30) |
| *Senecio vulgaris* | 2024 | LM | -21.26 | 20.68 | F(1,57) =<br>1.06 | 0.463 | R<sup>2</sup> = 0.02 (adj.<br>0.00) |
| *Stellaria media* | 2024 | LM | -6.94 | 37.96 | F(1,57) =<br>0.03 | 0.935 | R<sup>2</sup> = 0.00 (adj. -<br>0.02) |
| *Barbarea vulgaris* | 2025 | LM | 5.80 | 8.50 | F(1,58) =<br>0.47 | 0.661 | R<sup>2</sup> = 0.01 (adj. -<br>0.01) |
| *Centaurea jacea* | 2025 | LM | -1.77 | 5.69 | F(1,58) =<br>0.10 | 0.840 | R<sup>2</sup> = 0.00 (adj. -<br>0.02) |
| *Lychnis flos-cuculi* | 2025 | LM | 42.32 | 55.98 | F(1,54) =<br>0.57 | 0.622 | R<sup>2</sup> = 0.01 (adj. -<br>0.01) |
| *Plantago lanceolata* | 2025 | LM | 4.25 | 6.41 | F(1,54) =<br>0.44 | 0.667 | R<sup>2</sup> = 0.01 (adj. -<br>0.01) |
| *Plantago media* | 2025 | LM | 3.40 | 3.07 | F(1,58) =<br>1.23 | 0.450 | R<sup>2</sup> = 0.02 (adj.<br>0.00) |
| **Seed mass** |  |  |  |  |  |  |  |
| *Bupleurum rotundifolium* | 2024 | LM | -0.18 | 0.11 | F(1,55) =<br>2.45 | 0.262 | R<sup>2</sup> = 0.04 (adj.<br>0.03) |
| *Cardamine hirsuta* | 2024 | LM | -0.00 | 0.01 | F(1,52) =<br>0.00 | 0.978 | R<sup>2</sup> = 0.00 (adj. -<br>0.02) |
| *Plantago lanceolata* | 2024 | LM | -0.09 | 0.08 | F(1,49) =<br>1.12 | 0.454 | R<sup>2</sup> = 0.02 (adj.<br>0.00) |
| *Plantago media* | 2024 | LM | -0.05 | 0.02 | F(1,56) = 6.70 | **0.039** | R<sup>2</sup> = 0.11 (adj. 0.09) |
| *Senecio vulgaris* | 2024 | LM | 0.04 | 0.04 | F(1,54) = 1.00 | 0.475 | R<sup>2</sup> = 0.02 (adj. - 0.00) |
| *Stellaria media* | 2024 | LM | -0.12 | 0.10 | F(1,43) = 1.57 | 0.374 | R<sup>2</sup> = 0.04 (adj. 0.01) |
| *Barbarea vulgaris* | 2025 | LM | 0.05 | 0.05 | F(1,32) = 0.75 | 0.557 | R<sup>2</sup> = 0.02 (adj. - 0.01) |
| *Centaurea jacea* | 2025 | LM | 0.02 | 0.15 | F(1,43) = 0.01 | 0.964 | R<sup>2</sup> = 0.00 (adj. - 0.02) |
| *Lychnis flos-cuculi* | 2025 | LM | 0.01 | 0.01 | F(1,37) = 0.63 | 0.605 | R<sup>2</sup> = 0.02 (adj. - 0.01) |
| *Plantago lanceolata* | 2025 | LM | -0.06 | 0.13 | F(1,51) = 0.23 | 0.764 | R<sup>2</sup> = 0.00 (adj. - 0.02) |
| *Plantago media* | 2025 | LM | 0.04 | 0.10 | F(1,55) = 0.21 | 0.764 | R<sup>2</sup> = 0.00 (adj. - 0.01) |
| **Adult biomass** |  |  |  |  |  |  |  |
| *Barbarea vulgaris* | 2024 | LM | -2.09 | 1.99 | F(1,58) = 1.11 | 0.454 | R<sup>2</sup> = 0.02 (adj. 0.00) |
| *Bupleurum rotundifolium* | 2024 | LM | -0.78 | 0.52 | F(1,56) = 2.25 | 0.281 | R<sup>2</sup> = 0.04 (adj. 0.02) |
| *Cardamine hirsuta* | 2024 | LM | -0.54 | 0.45 | F(1,58) = 1.45 | 0.394 | R<sup>2</sup> = 0.02 (adj. 0.01) |
| *Centaurea jacea* | 2024 | LM | -3.35 | 1.51 | F(1,58) = 4.90 | 0.086 | R<sup>2</sup> = 0.08 (adj. 0.06) |
| *Lychnis flos-cuculi* | 2024 | LM | -0.69 | 0.55 | F(1,58) = 1.60 | 0.371 | R<sup>2</sup> = 0.03 (adj. 0.01) |
| *Plantago lanceolata* | 2024 | LM | -11.70 | 2.22 | F(1,57) = 27.81 | **< 0.001** | R<sup>2</sup> = 0.33 (adj. 0.32) |
| *Plantago media* | 2024 | LM | -5.99 | 0.83 | F(1,58) = 52.74 | **< 0.001** | R<sup>2</sup> = 0.48 (adj. 0.47) |
| *Senecio vulgaris* | 2024 | LM | -1.01 | 0.73 | F(1,58) = 1.93 | 0.320 | R<sup>2</sup> = 0.03 (adj. 0.02) |
| *Stellaria media* | 2024 | LM | -1.40 | 0.49 | F(1,58) = 7.97 | **0.024** | R<sup>2</sup> = 0.12 (adj. 0.11) |
| *Barbarea vulgaris* | 2025 | LM | 1.95 | 1.80 | F(1,57) = 1.18 | 0.454 | R<sup>2</sup> = 0.02 (adj. 0.00) |
| *Centaurea jacea* | 2025 | LM | -0.23 | 2.55 | F(1,59) = 0.01 | 0.964 | R<sup>2</sup> = 0.00 (adj. - 0.02) |
| *Lychnis flos-cuculi* | 2025 | LM | 2.13 | 2.93 | F(1,54) = 0.53 | 0.634 | R<sup>2</sup> = 0.01 (adj. - 0.01) |
| *Plantago lanceolata* | 2025 | LM | 1.38 | 2.46 | F(1,54) = 0.31 | 0.731 | R<sup>2</sup> = 0.01 (adj. - 0.01) |
| *Plantago media* | 2025 | LM | 7.11 | 2.14 | F(1,58) = 10.98 | **0.007** | R<sup>2</sup> = 0.16 (adj. 0.14) |

| Species | Year | Model | Estimate | SE | Test Stat | P-value<br>(adjusted) | Effect Size* |
| --- | --- | --- | --- | --- | --- | --- | --- |

Note: Models: LM = Linear Model, LMM = Linear Mixed Model, GLM = Generalized Linear Model \* R<sup>2</sup>c = conditional R<sup>2</sup> (fixed + random effects)

## Notes

### Competing Interest Statement

The authors have declared no competing interest.

